# Interplay Between Cholesterol Concentration and Membrane Curvature in Liposomes Revealed by Molecular Dynamics Simulations

**DOI:** 10.64898/2026.07.15.738831

**Authors:** Ehsan Khodadadi, Mortaza Derakhshani-Molayousefi, Ehsaneh Khodadadi, Mahmoud Moradi

## Abstract

Liposomes are widely used as model membranes and nanoscale drug delivery systems, where cholesterol plays a key role in regulating bilayer structure and dynamics. However, how cholesterol concentration influences the structure and dynamics of liposome and how this influence is dependent on membrane curvature are not fully understood at the molecular level. In this work, coarse-grained molecular dynamics simulations using the MARTINI force field were employed to examine the concentration-dependent behavior of cholesterol in planar and curved membranes composed of cholesterol and unsaturated phospholipids, namely DOPC. More specifically, a planar lipid bilayer and an approximately 50-nm liposome were simulated to represent two extreme limits of small and large curvature, respectively. Increasing cholesterol concentration led to thicker membranes and reduced solvent exposure, consistent with cholesterol’s condensing effect. Membrane curvature enhanced interleaflet coupling and increased tail interdigitation relative to planar systems. Notably, DOPC flip-flop rate in spherical bilayers exhibited a non-monotonic dependence on cholesterol content, reflecting a balance between curvature-induced packing stress and cholesterol-driven ordering. These findings provide molecular-level insight into how cholesterol and curvature together shape the structure and dynamics of unsaturated lipid bilayers.

## 1. Introduction

Molecular dynamics (MD) simulations provide a powerful framework for probing the structure and dynamics of biological membranes at molecular and mesoscopic scales. By resolving lipid motions and interactions over time, MD simulations offer insight into the thermodynamic and kinetic factors that govern membrane organization and function [1, 2, 3]. Lipid bilayers are of particular interest because their structural and mechanical properties play a central role in cellular organization, signaling, and transport.

Cholesterol is a major component of eukaryotic membranes and strongly modulates bilayer thickness, fluidity, permeability, and mechanical rigidity through its interactions with neighboring phospholipids [4, 5]. While cholesterol can promote more ordered lipid arrangements under certain conditions, the extent to which such ordering translates into stable, spatially segregated domains in biological membranes remains an open question. Capturing cholesterol redistribution and transbilayer motion typically requires long simulation times, which are often inaccessible to all-atom MD approaches [6, 7]. Coarse-Grained (CG) MD models address these limitations by reducing atomic detail while retaining essential intermolecular interactions, enabling simulations of larger membrane systems over extended timescales. The MARTINI force field has been widely applied to lipid membranes and vesicles, allowing systematic investigation of curvature effects, lipid packing, and cholesterol-dependent membrane behavior [8, 9, 10]. Although CG models accelerate molecular dynamics relative to experiment, they provide valuable mechanistic insight into membrane organization that complements experimental observations.

From an applied perspective, liposomes spherical vesicles composed of one or more lipid bilayers are well-established carriers for drug delivery, where membrane composition critically influences stability and performance [11, 12]. Among commonly used phospholipids, 1,2-dioleoyl-sn-glycero-3phosphocholine (DOPC) is frequently employed due to its ability to form stable, fluid bilayers and its presence in several clinically approved liposomal formulations [13, 14, 15]. Cholesterol is routinely incorporated into such systems to enhance membrane stability, reduce leakage, and improve therapeutic efficacy [16, 17].

Experimental studies on saturated phosphatidylcholine systems often report reproducible macroscopic behavior near specific lipid-to-cholesterol ratios, such as 70:30, under controlled preparation conditions [18]. However, these formulation benchmarks reflect long-term stability and bulk properties rather than the molecular-scale mechanisms governing membrane organization. In contrast, unsaturated lipids such as DOPC exhibit distinct packing behavior, elasticity, and cholesterol interactions, particularly in curved membrane geometries [13, 19]. Membrane curvature introduces packing frustration and leaflet asymmetry, which can influence lipid organization and transbilayer exchange in a geometry-dependent manner [20].

In this work, we use CG MARTINI simulations to systematically examine how cholesterol concentration modulates the structural and dynamic properties of DOPC membranes in planar bilayers and spherical vesicles. We focus on cholesterol-dependent trends in membrane thickness, solvent-accessible surface area (SASA), lipid tail interdigitation, segmental order parameters, and transbilayer lipid exchange. Rather than proposing an optimal formulation, our goal is to elucidate how cholesterol and membrane curvature jointly shape the molecular organization of unsaturated lipid bilayers. By emphasizing relative trends and mechanistic insight, this study provides a framework for interpreting cholesterol’s role in curved membranes at the nanoscale and complements existing experimental and computational work.

## 2. Materials and Methods

CG-MD simulations were performed to investigate the effects of cholesterol concentration and membrane curvature on the structural and dynamic properties of DOPC–cholesterol membranes. Both planar bilayers and spherical vesicles were considered to enable direct comparison between flat and curved geometries under otherwise identical simulation conditions. All simulations employed the MARTINI 2.2 CG force field [21, 22], which has been extensively validated for lipid–cholesterol systems. Although newer MARTINI parameterizations are available, the present study focuses on relative, composition-dependent trends within a consistent force-field framework rather than on quantitative agreement with experimental observables [23, 8].

### 2.1. Simulation Details

Planar bilayers were constructed using the CHARMM-GUI Martini Bilayer Maker, while spherical vesicles were generated with the CHARMM-GUI Martini Vesicle Maker [24, 25, 26]. These tools provided fully parameterized input files compatible with GROMACS (version 2024), which was used for all simulations [27]. Five lipid compositions were examined, with DOPC-to-cholesterol molar ratios of 100:0, 90:10, 80:20, 70:30, and 60:40. This range spans cholesterol fractions commonly employed in computational and experimental studies to probe cholesterol-dependent membrane behavior [28].

For planar systems, simulations were performed in rectangular boxes with dimensions of 300 Å × 300 Å × 85 Å, providing a water thickness of approximately 22.5 Å on each side of the bilayer. Spherical systems were simulated in cubic boxes of 640 Å × 640 Å × 640 Å, enclosing vesicles with an outer radius of approximately 260 Å. During vesicle construction, a transient cylindrical water pore was introduced to facilitate bilayer assembly; this pore was removed prior to production simulations, and all analyses were performed only after complete membrane resealing was confirmed.

All systems were solvated using MARTINI CG water beads, and NaCl was added to a concentration of 0.15 M to mimic physiological ionic strength. Energy minimization was carried out using the steepest descent algorithm for 5,000 steps to remove unfavorable steric contacts [29]. Systems were subsequently equilibrated under NVT conditions to stabilize temperature, followed by NPT equilibration to relax pressure and density. A time step of 20 fs was used throughout, consistent with standard MARTINI simulation protocols. Temperature was maintained at 310 K using the velocity-rescaling thermostat with a stochastic term [30, 31]. During equilibration, pressure coupling was performed using the Berendsen barostat [32], followed by the Parrinello–Rahman barostat during production runs [33].

Production simulations were carried out for 7 *µ*s for each system, with three independent replicas per composition and geometry, yielding a total of 30 simulations. Owing to the accelerated dynamics inherent to the MARTINI framework, simulated timescales were interpreted in a relative sense rather than as direct representations of experimental kinetics.

### 2.2. Analysis

Trajectory analysis was performed using in-house Python scripts based on MDAnalysis [34, 35], with additional support from NumPy [36], IPython [37], and Matplotlib [38]. Visualization was carried out using VMD [39]. GROMACS analysis tools were also employed for selected calculations, including SASA [40]. Structural and dynamic metrics were evaluated after equilibration to quantify cholesterol and curvature dependent trends in membrane behavior [41, 42].

Membrane structural and dynamic properties were analyzed separately for planar bilayers and spherical vesicles to account explicitly for differences in geometry and membrane curvature.

For planar bilayers, all quantities were evaluated over the entire membrane following equilibration. For spherical bilayers, lipid properties were analyzed over the full vesicle surface unless otherwise noted. No spatial subdivision of the vesicle surface was applied; all reported quantities therefore represent global averages over all lipids, capturing curvature-induced heterogeneity in a statistical manner.

The analyzed properties included membrane thickness, lipid interdigitation, SASA, segmental order parameters (*S_CD_*), lipid transbilayer exchange, and vesicle radius probability density distributions, as described below.

Unless otherwise stated, all analyses were performed on equilibrated segments of the production trajectories. Equilibration was assessed by monitoring the convergence of key observables, including membrane thickness and lipid tail order parameters. Only trajectory segments exhibiting no systematic drift in these quantities were included in the analysis.

Statistical uncertainties were estimated using block averaging over equilibrated trajectory segments, with block lengths chosen to ensure convergence of the standard error. Reported values therefore represent time-averaged properties with associated uncertainty estimates. For dynamic observables such as lipid flip-flopping, additional persistence criteria were applied to exclude transient fluctuations and ensure that only sustained events were counted.

#### 2.2.1. Membrane Thickness

Membrane thickness provides a global structural measure of bilayer organization and lipid packing. For planar bilayers, thickness was defined as the absolute separation between the mean positions of the phospholipid headgroup beads in the two opposing leaflets along the membrane normal:

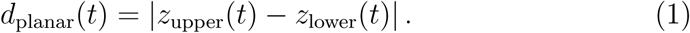

This definition yields a robust global thickness metric suitable for relative comparisons across cholesterol concentrations.

For spherical vesicles, a global radial definition was employed. At each frame, the vesicle center was defined as the center of mass of all membrane lipids, excluding solvent and ions. Headgroup beads were assigned to inner and outer leaflets, and thickness was calculated as

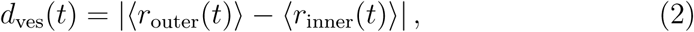

where *r* denotes the distance from the vesicle center and ⟨·⟩ indicates averaging over lipids in each leaflet. The reported thickness values therefore represent global averages rather than local curvature-resolved measurements.

This approach enables consistent comparison between planar and curved membranes while focusing on cholesterol-dependent trends rather than absolute experimental thickness values [43, 44].

#### 2.2.2. Lipid Interdigitation

Lipid interdigitation quantifies the extent of tail overlap between opposing leaflets and provides insight into interleaflet packing and geometric constraints. For planar bilayers, interdigitation was computed from the overlap of lipid tail bead distributions along the global membrane normal relative to the bilayer midplane, following established methods [45].

For spherical bilayers, where a single global membrane normal is not defined, interdigitation was evaluated using a radial definition relative to the vesicle center. Lipid tail positions were referenced to the vesicle midplane, yielding a global measure of tail overlap appropriate for curved membranes [46].

Interdigitation values were interpreted as geometric indicators of curvature-induced packing effects rather than direct measures of membrane stability or mechanical strength. Consistent with previous studies, spherical bilayers exhibited higher interdigitation than planar systems due to curvature-induced constraints on lipid packing [47].

#### 2.2.3. Solvent-Accessible Surface Area

The SASA was calculated to characterize lipid exposure to the aqueous environment and changes in molecular packing. SASA calculations were performed using the gmx sasa tool in GROMACS, which implements the Shrake–Rupley algorithm [48]. A fixed probe radius was used consistently across all systems.

SASA was computed separately for DOPC, cholesterol, and the combined lipid population on a per lipid basis and averaged over equilibrated trajectory segments. For spherical vesicles, SASA values represent global averages over the entire vesicle surface. Accordingly, SASA is interpreted here as a qualitative structural descriptor of lipid organization rather than as a direct measure of membrane permeability or stability [49, 50].

#### 2.2.4. Segmental Order Parameter

Lipid tail orientational order was quantified using the *S_CD_*, defined as

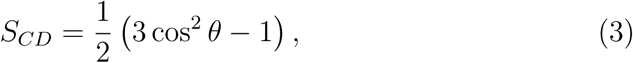

where *θ* is the angle between a lipid tail bond vector and the local membrane normal[51, 52].

In the MARTINI CG representation, *S_CD_* was computed using bond vectors along the lipid acyl chains, providing an effective measure of tail ordering suitable for relative comparisons across systems [53, 54]. For planar bilayers, the membrane normal was defined along the global bilayer normal. For spherical bilayers, lipid orientations were evaluated relative to locally defined membrane normals based on the instantaneous geometry of nearby headgroups, following curvature aware approaches reported previously [19, 55].

Order parameters were calculated separately for the sn-1 and sn-2 chains and averaged over all lipids and analyzed frames. The resulting values reflect global lipid tail ordering rather than local mechanical properties.

#### 2.2.5. Lipid Transbilayer Exchange

Lipid transbilayer exchange (flip-flopping) was quantified to characterize relative translocation dynamics of DOPC and cholesterol molecules. Lipids were assigned to leaflets on a frame-by-frame basis using a geometric criterion based on headgroup positions relative to the instantaneous bilayer midplane [56, 19].

A flip-flop event was defined as a transition between opposing leaflets that persisted for at least 50 ns. This persistence criterion was applied to exclude transient midplane crossings and short-lived fluctuations. Flip-flop frequencies were computed as the number of sustained transitions per lipid per microsecond and averaged over independent replicas.

Due to the accelerated dynamics inherent to the MARTINI framework, flip-flop frequencies are reported as relative, model-dependent measures intended for comparison across cholesterol concentrations and membrane geometries rather than as quantitative estimates of experimental rates.

#### 2.2.6. Cholesterol-Dependent Vesicle Radius Distributions

Vesicle geometry was characterized by computing probability density distributions of the vesicle radius as a function of cholesterol concentration. At each frame, the radius was determined from the distances of membrane headgroup beads to the vesicle center of mass, defined as in the thickness analysis.

Because lipid number and simulation conditions were held constant, shifts in radius distributions reflect cholesterol-dependent relaxation and packing effects within the CG model rather than externally imposed size changes [57, 19]. These distributions are therefore interpreted as comparative geometric descriptors rather than quantitative predictors of experimental vesicle sizes [58].

## 3. Results and Discussion

### 3.1. Structural Responses of Lipid Bilayers: Thickness and Interdigitation

Membrane thickness and lipid tail interdigitation are key structural descriptors that reflect lipid packing, tail ordering, and interleaflet organization in bilayer membranes. These properties influence how membranes accommodate molecular composition and curvature and are therefore particularly relevant for vesicular systems such as liposomes. MD simulations have been widely used to examine how cholesterol alters these structural features across different lipid compositions and membrane geometries [59, 60].

Using CG MARTINI simulations, we examined DOPC/cholesterol membranes in both planar bilayers and spherical vesicles over a range of cholesterol concentrations (Fig. 1). In both geometries, increasing cholesterol concentration leads to a systematic increase in membrane thickness. This behavior is consistent with cholesterol’s well established condensing effect, whereby sterol–lipid interactions promote straighter lipid tails and tighter hydrocarbon packing [43, 61]. The monotonic thickening observed here indicates that cholesterol driven tail ordering is preserved in both flat and curved DOPC membranes within the CG framework.

**Figure 1:**
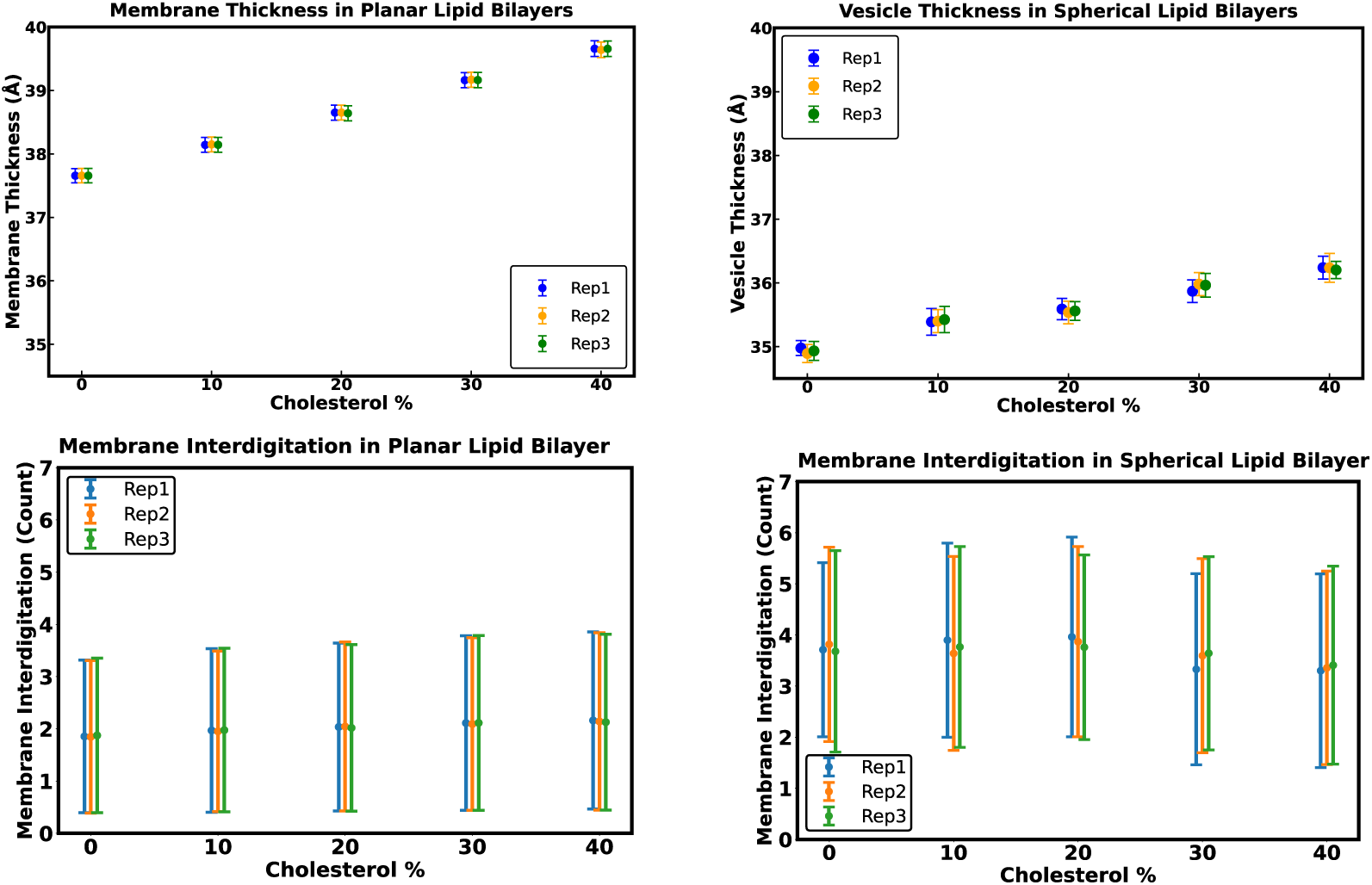
Cholesterol-dependent membrane thickness and lipid tail interdigitation in planar and spherical bilayers. Top: Membrane thickness increases with cholesterol concentration in both planar (left) and spherical (right) bilayers, with greater variability in the curved geometry. **Bottom:** Lipid tail interdigitation shows little dependence on cholesterol in planar bilayers (left) but is consistently higher in spherical bilayers (right), reflecting curvature-induced packing constraints.

In contrast to membrane thickness, lipid tail interdigitation exhibits a strong dependence on membrane geometry and only a weak dependence on cholesterol concentration. Planar bilayers show nearly constant interdigitation values across all cholesterol fractions, indicating that cholesterol induced ordering primarily affects intraleaflet packing rather than interleaflet tail overlap in flat membranes. Spherical bilayers, however, consistently display higher interdigitation than planar systems at all compositions. This enhancement arises from curvature induced packing constraints that promote greater tail overlap as the membrane accommodates differences in surface area between the inner and outer leaflets.

These observations demonstrate that cholesterol and curvature influence bilayer structure through distinct mechanisms. Cholesterol primarily modulates lipid ordering and bilayer thickness, whereas curvature governs the extent of interleaflet tail overlap. Importantly, neither thickness nor interdigitation should be interpreted as direct measures of membrane stability or vesicle robustness; instead, they serve as structural descriptors that highlight how lipid packing responds to compositional and geometric constraints.

#### 3.1.1. Membrane Thickness

Membrane thickness provides a global measure of bilayer organization and reflects changes in lipid tail ordering and packing density [62, 63]. In planar DOPC/cholesterol bilayers (Fig. 1, top left), membrane thickness increases monotonically with cholesterol concentration. The response is approximately linear over the investigated range, and the three independent replicas show minimal variability, indicating a uniform packing response in the absence of curvature.

Spherical bilayers exhibit the same qualitative cholesterol dependent thickening trend (Fig. 1, top right). However, the magnitude of the thickness increase is reduced relative to planar membranes, and replicate-to-replicate variability is more pronounced. This increased dispersion reflects curvature-induced packing heterogeneity: in vesicles, the inner and outer leaflets experience different surface areas and lateral stresses, leading to less uniform lipid alignment than in flat bilayers [61, 44].

These results indicate that cholesterol-induced thickening is a robust feature of DOPC membranes, but its expression is modulated by membrane curvature. Curved geometries partially frustrate uniform tail alignment, resulting in increased variability of global thickness measurements without altering the overall direction of the cholesterol-dependent trend.

#### 3.1.2. Interdigitation

Lipid tail interdigitation quantifies the degree of overlap between tails from opposing leaflets and serves as a geometric descriptor of interleaflet packing. In planar DOPC/cholesterol bilayers (Fig. 1, bottom left), interdigitation values remain nearly invariant across cholesterol concentrations. This weak dependence indicates that cholesterol does not substantially alter tail overlap in flat membranes, consistent with previous simulations showing that cholesterol primarily reorganizes intraleaflet packing rather than promoting interleaflet mixing [64, 65].

In spherical bilayers (Fig. 1, bottom right), interdigitation values are systematically higher than in planar systems but show only modest sensitivity to cholesterol concentration. The elevated interdigitation arises from geometric constraints imposed by curvature: the inner leaflet must accommodate a smaller surface area than the outer leaflet, leading to differential tail orientation and enhanced apparent overlap between leaflets. Similar curvature-driven increases in interdigitation have been reported in previous studies of vesicular and highly curved membrane systems [47, 46].

Together, these findings indicate that membrane curvature, rather than cholesterol concentration, is the dominant factor controlling interleaflet tail overlap in DOPC bilayers within the CG framework.

#### 3.1.3. Integrated View of Cholesterol–Curvature Interplay

The combined analysis of membrane thickness and interdigitation highlights that cholesterol content and membrane curvature affect bilayer structure through complementary but largely decoupled mechanisms (Fig. 1). In planar bilayers, cholesterol produces the expected monotonic increase in thickness while leaving interdigitation essentially unchanged. In spherical bilayers, curvature introduces geometric constraints that elevate interdigitation and increase variability in thickness, even as the overall cholesterol-dependent thickening trend is preserved.

These results underscore that structural trends derived from planar bilayer simulations cannot be directly transferred to curved membrane systems. In vesicles, curvature-induced packing constraints can dominate over compositional effects for certain structural descriptors, leading to broader distributions of global properties. Accordingly, the trends reported here should be interpreted as geometry-dependent responses within a CG framework rather than as direct predictors of vesicle stability or experimental membrane behavior.

### 3.2. Solvent-Accessible Surface Area

Figure 2 summarizes the cholesterol-dependent behavior of SASA in planar and spherical DOPC bilayers. Here, SASA is used as a qualitative structural descriptor of lipid exposure to the aqueous environment and reflects changes in molecular packing associated with membrane composition and geometry.

**Figure 2:**
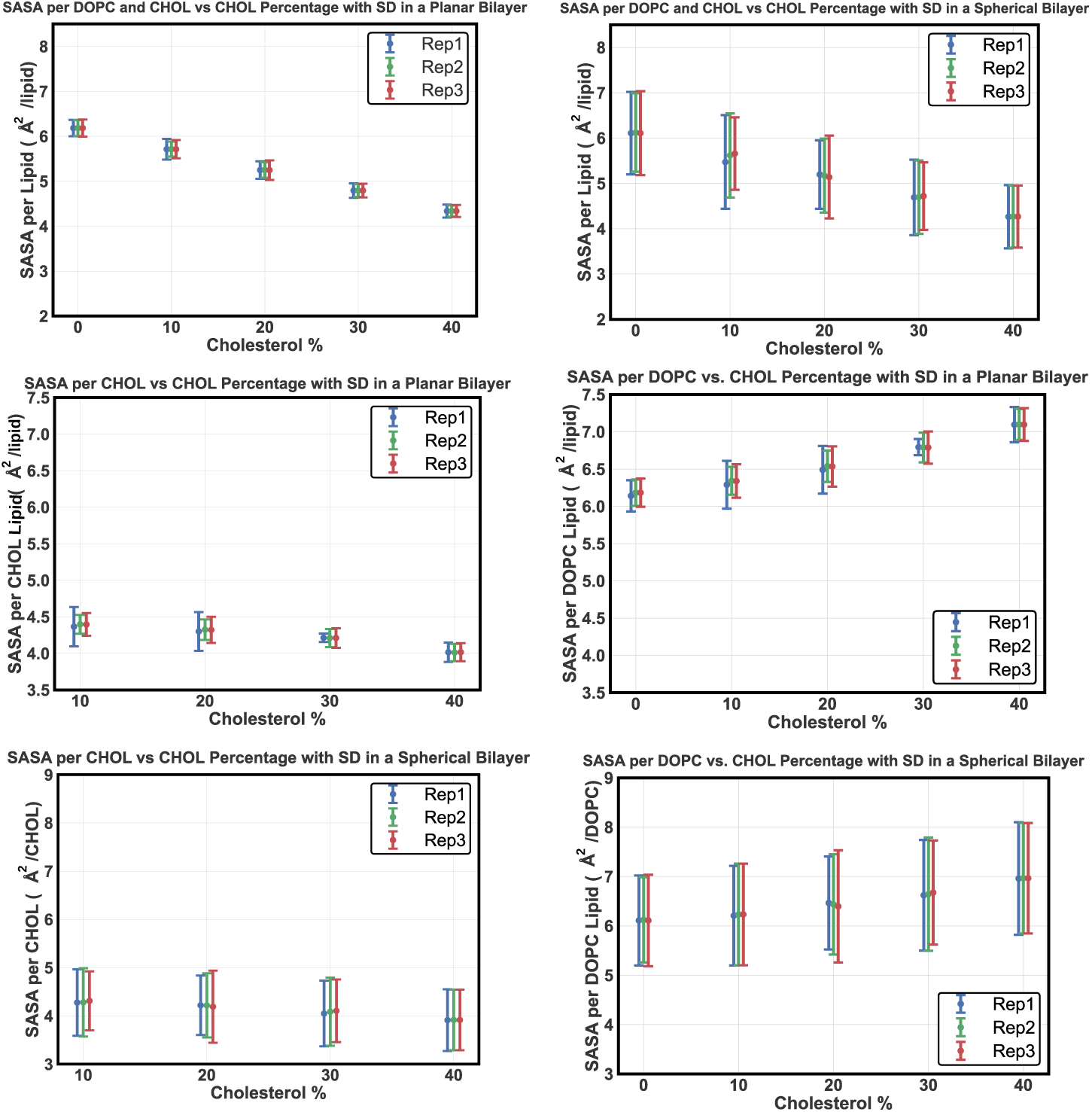
Cholesterol-dependent solvent-accessible surface area (SASA) in planar and spherical bilayers. Top Left: SASA per lipid (DOPC + CHOL) in planar bilayers decreases monotonically with increasing cholesterol concentration, consistent with cholesterol-induced condensation. **Top Right:** SASA per lipid in spherical bilayers follows the same qualitative trend, with increased replica-to-replica variability reflecting curvature-induced packing heterogeneity. **Middle Left:** SASA per cholesterol molecule in planar bilayers decreases with cholesterol fraction, indicating reduced solvent exposure at higher sterol content. **Middle Right:** SASA per DOPC molecule in planar bilayers shows a modest increase with cholesterol concentration due to compositional normalization effects. **Bottom Left:** SASA per cholesterol molecule in spherical bilayers exhibits a comparable decreasing trend with broader distributions. **Bottom Right:** SASA per DOPC molecule in spherical bilayers displays cholesterol-dependent behavior with enhanced variability associated with membrane curvature.

In planar bilayers (Fig. 2, top left), the SASA per lipid (DOPC + CHOL) decreases smoothly and monotonically with increasing cholesterol concentration. This trend is consistent with the well-established condensing effect of cholesterol, whereby tighter lipid packing reduces overall solvent exposure. The small replica-to-replica variation indicates a relatively uniform lateral organization in flat membranes [66, 49, 50].

Spherical bilayers (Fig. 2, top right) exhibit the same qualitative decrease in SASA per lipid with increasing cholesterol content, but with noticeably larger variability between replicas. This increased spread reflects curvature-induced heterogeneity in local packing environments rather than the formation of discrete domains. In curved vesicles, lipids experience continuously varying geometric constraints, leading to broader distributions in SASA-based observables [67, 68].

Cholesterol-specific SASA values decrease with increasing cholesterol fraction in both planar and spherical systems (Fig. 2, middle left and bottom left), consistent with increased sterol–lipid and sterol–sterol contacts that reduce solvent exposure. As for total SASA, spherical bilayers show broader distributions, attributable to curvature-dependent packing variability rather than qualitatively different cholesterol behavior.

In contrast, the SASA per DOPC molecule increases modestly with cholesterol concentration in planar bilayers (Fig. 2, middle right). This increase reflects a compositional normalization effect: as cholesterol replaces DOPC, the remaining phospholipids experience a higher average solvent exposure per molecule rather than an expansion of individual lipid footprints. A similar tendency is observed in spherical bilayers (Fig. 2, bottom right), albeit with larger fluctuations arising from curvature, leaflet asymmetry, and heterogeneous local environments [68, 69].

Overall, the SASA analysis demonstrates that cholesterol reduces solvent exposure at the bilayer level in both planar and curved DOPC membranes, while membrane curvature primarily increases the variability of SASA measurements without altering the qualitative cholesterol-dependent trends. These results should therefore be interpreted as structural signatures of lipid packing and geometry within a CG framework, rather than as direct predictors of membrane stability or liposome performance in experimental systems [70].

### 3.3. Lipid Tail Ordering

The segmental order parameter (*S_CD_*) was used to quantify lipid tail orientational ordering and to assess how cholesterol and membrane geometry influence chain organization in DOPC bilayers. Higher *S_CD_* values correspond to increased alignment of lipid tails along the membrane normal and reduced conformational flexibility [53, 71].

In planar bilayers (Fig. 3, top panels), both the sn-1 and sn-2 tails of DOPC exhibit a clear, monotonic increase in *S_CD_*with increasing cholesterol concentration. This trend reflects the well-established ordering effect of cholesterol, which restricts lipid tail motion and promotes more extended chain conformations [72, 54]. The increase in *S_CD_*is smooth and highly reproducible across independent replicas, consistent with the relatively uniform packing environment of flat membranes.

**Figure 3:**
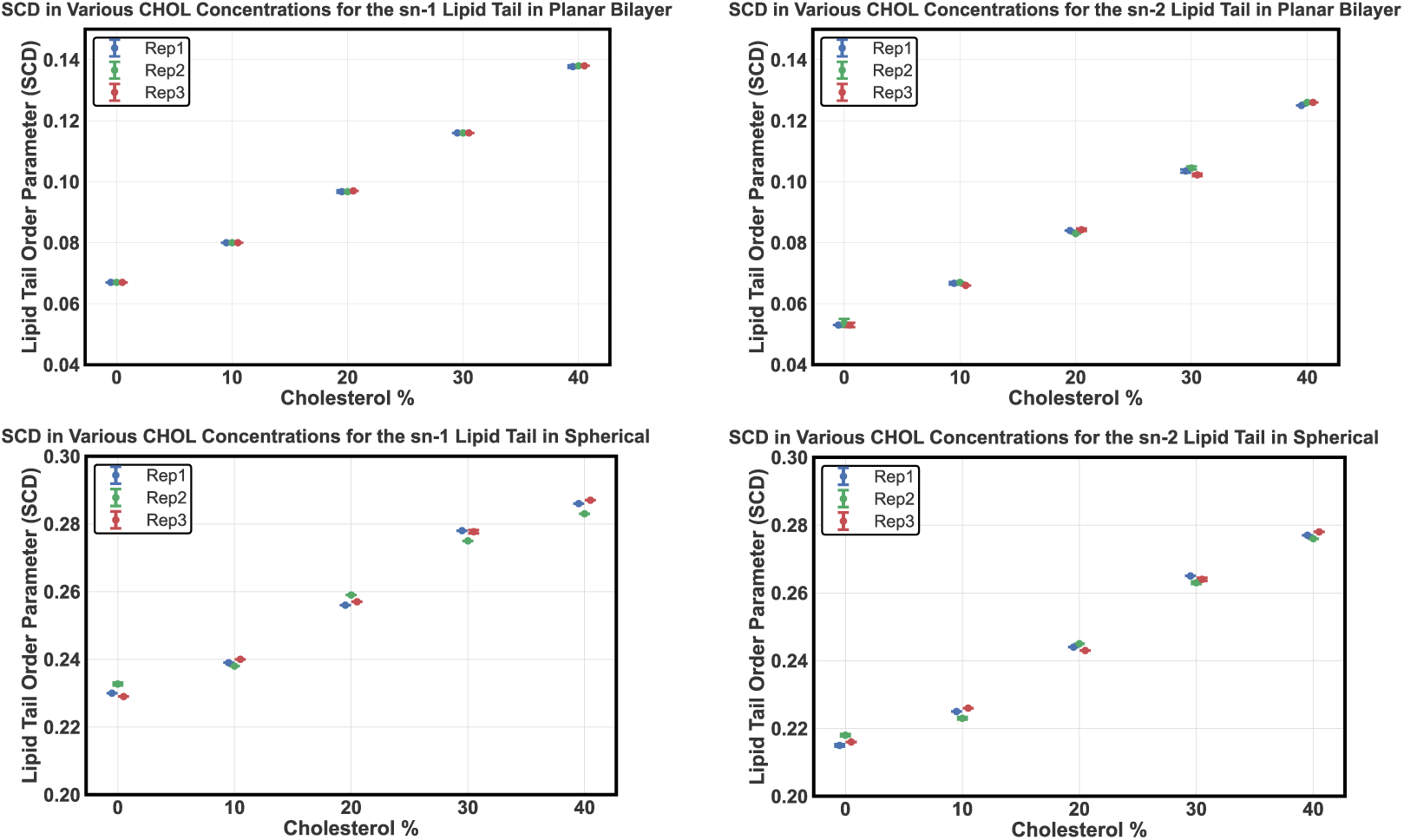
Cholesterol Dependence of Lipid Tail Ordering in Planar and Spherical Bilayers. Top panels: *S_CD_* for the sn-1 (left) and sn-2 (right) DOPC tails in planar bilayers, showing a monotonic increase with cholesterol concentration. **Bottom panels:** Corresponding *S_CD_* profiles in spherical bilayers. Ordering increases with cholesterol as in planar systems, with systematically higher baseline values reflecting curvature-related geometric constraints.

Spherical bilayers display the same qualitative cholesterol-dependent increase in *S_CD_* for both sn-1 and sn-2 tails (Fig. 3, bottom panels). At a given cholesterol concentration, *S_CD_* values are systematically higher than those observed in planar bilayers, including at zero cholesterol. This offset reflects curvature induced geometric constraints on lipid packing and tail orientation in the vesicle geometry rather than a direct enhancement of molecular order. Importantly, membrane curvature does not alter the overall cholesterol response: lipid tail ordering increases monotonically with cholesterol concentration in both planar and spherical bilayers. Replica-to-replica variability is modestly larger in spherical systems, consistent with the greater structural heterogeneity inherent to curved membranes.

Overall, the *S_CD_*analysis demonstrates that cholesterol enhances lipid tail ordering in DOPC bilayers independently of membrane geometry, while curvature primarily shifts the baseline level of apparent ordering within the CG framework.

### 3.4. Flip-Flop Dynamics

Lipid flip-flopping, defined as the transbilayer migration of lipid molecules between opposing leaflets, provides insight into membrane dynamics, packing constraints, and compositional relaxation in multicomponent bilayers. Figures 4 and 5 summarize the cholesterol- and geometry-dependent flip-flop behavior of cholesterol and DOPC in planar and spherical DOPC/cholesterol bilayers within the CG MARTINI framework.

**Figure 4:**
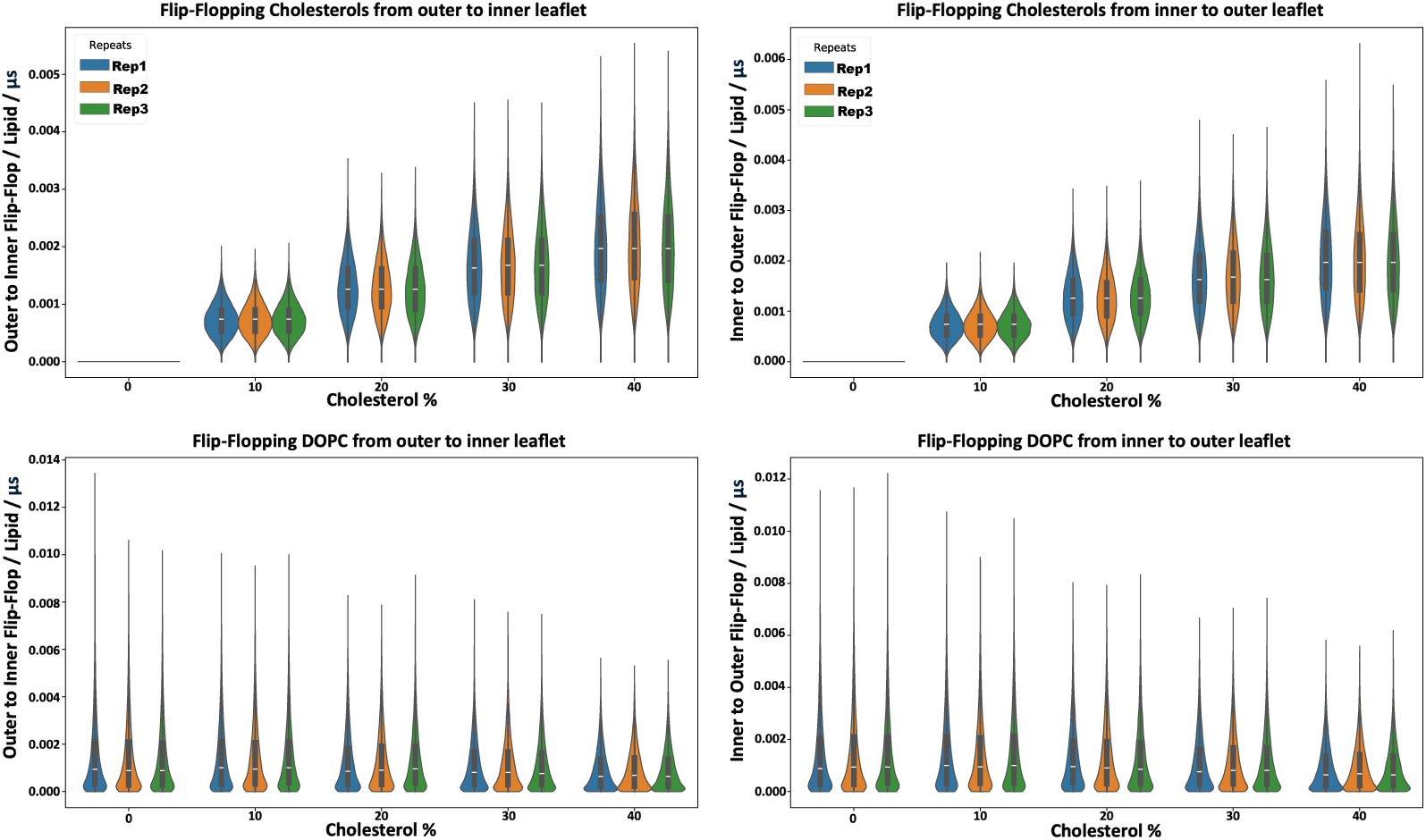
Cholesterol-dependent flip-flop dynamics of DOPC and cholesterol in planar bilayers. Violin plots show flip-flop frequency distributions from three independent replicas at different cholesterol concentrations. **Top Left and Top Right:** Cholesterol flip-flop increases monotonically with cholesterol concentration in both outer-to-inner and inner-to-outer directions, consistent with enhanced sterol translocation in the CG model. **Bottom Left and Bottom Right:** DOPC flip-flop remains low across all compositions, with only minor variations and a slight reduction at higher cholesterol fractions, indicating limited phospholipid translocation on the simulated timescales.

**Figure 5:**
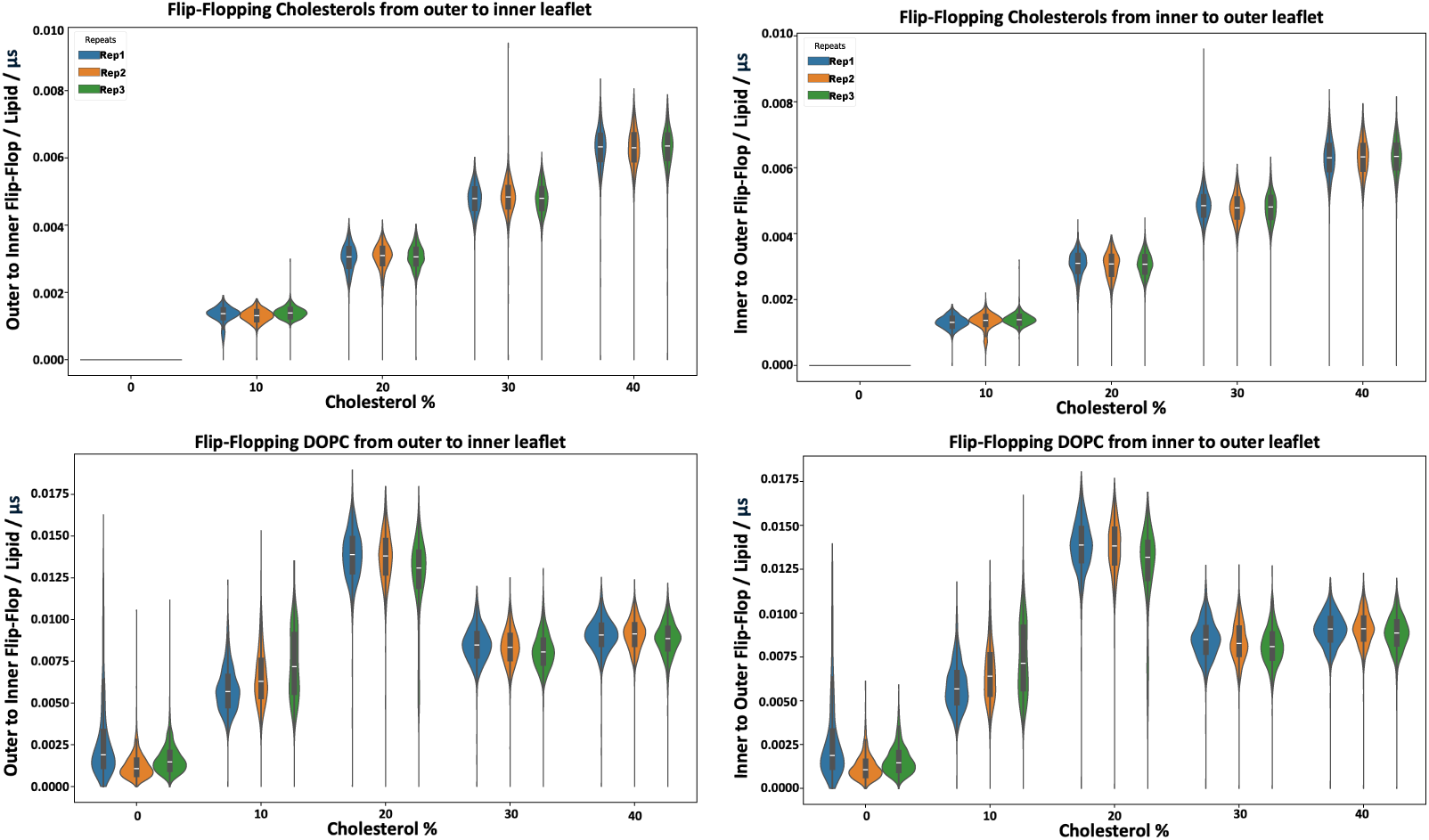
Cholesterol-dependent flip-flop behavior in spherical lipid bilayers. Violin plots show flip-flop frequency distributions from three independent replicas of spherical DOPC/CHOL bilayers. **Top Left and Top Right:** Cholesterol flip-flop increases with cholesterol concentration and reaches its highest values at 40%, consistent with enhanced sterol translocation in curved membranes within the CG framework. **Bottom Left and Bottom Right:** DOPC flip-flop displays a non-monotonic dependence on cholesterol concentration, with a maximum near 20%, reflecting a crossover between curvature-induced leaflet stress and cholesterol-induced ordering in the vesicle geometry.

In planar bilayers (Fig. 4), cholesterol exhibits a clear and monotonic increase in flip-flop frequency with increasing cholesterol concentration in both outer-to-inner and inner-to-outer directions. This behavior is consistent with cholesterol’s small polar headgroup and rigid sterol core, which reduce the effective energetic barrier for transbilayer migration compared with phospholipids in CG models [73, 74]. Transient sterol reorientation and interfacial interactions facilitate repeated leaflet crossings as cholesterol content increases, leading to enhanced sterol mobility across the bilayer [75, 76].

In contrast, DOPC flip-flopping in planar bilayers remains low across all cholesterol concentrations, with only modest variation and a slight decrease at higher cholesterol fractions. This behavior reflects the large polar headgroup of phosphatidylcholine lipids and the absence of strong driving forces for sustained transbilayer exchange in flat membranes. The observed DOPC flip-flop events therefore represent rare, model-dependent fluctuations within the MARTINI framework rather than quantitative predictions of experimentally relevant flip-flop rates, which typically occur on much longer timescales [56].

In spherical bilayers (Fig. 5), cholesterol flip-flopping is further enhanced and shows a stronger dependence on cholesterol concentration than in planar systems, reaching its highest values at 40% cholesterol. This enhancement is consistent with curvature-induced packing frustration and local density variations in vesicles, which lower the effective barrier for sterol reorientation and promote transbilayer migration in curved membranes [77].

DOPC flip-flopping in spherical bilayers exhibits a non-monotonic dependence on cholesterol concentration, with the highest flip-flop rates observed near 20%. Rather than indicating a universal optimal cholesterol fraction, this behavior reflects a crossover regime in which curvature-induced leaflet stress promotes phospholipid translocation, while cholesterol-driven ordering progressively suppresses lipid mobility at higher concentrations. At elevated cholesterol levels, increased chain ordering and reduced lateral diffusion limit DOPC transbilayer exchange, resulting in lower flip-flop frequencies [56, 77]. Overall, these results demonstrate that cholesterol flip-flopping is strongly enhanced by both increasing cholesterol concentration and membrane curvature, whereas DOPC flip-flopping remains limited and highly sensitive to the balance between curvature effects and cholesterol-induced ordering. Importantly, the reported flip-flop frequencies should be interpreted as qualitative, comparative measures within a CG framework rather than as direct estimates of experimental transbilayer transport rates.

### 3.5. Vesicle Radius Distributions

Probability density distributions of the vesicle radius provide a global geometric descriptor of how spherical lipid bilayers relax as a function of cholesterol concentration under fixed simulation conditions. Changes in vesicle radius within CG models reflect composition-dependent lipid packing and membrane organization rather than direct control of vesicle size [78, 63].

Figure 6 shows that both the inner and outer radius distributions shift systematically toward smaller values as cholesterol concentration increases from 0% to 40%. Because lipid number and simulation parameters are held constant, this trend indicates that higher cholesterol fractions promote slightly more compact vesicle configurations within the present CG framework. Such behavior is consistent with cholesterol’s condensing effect, which reduces the effective area per lipid and enhances bilayer order [56, 19, 79].

**Figure 6:**
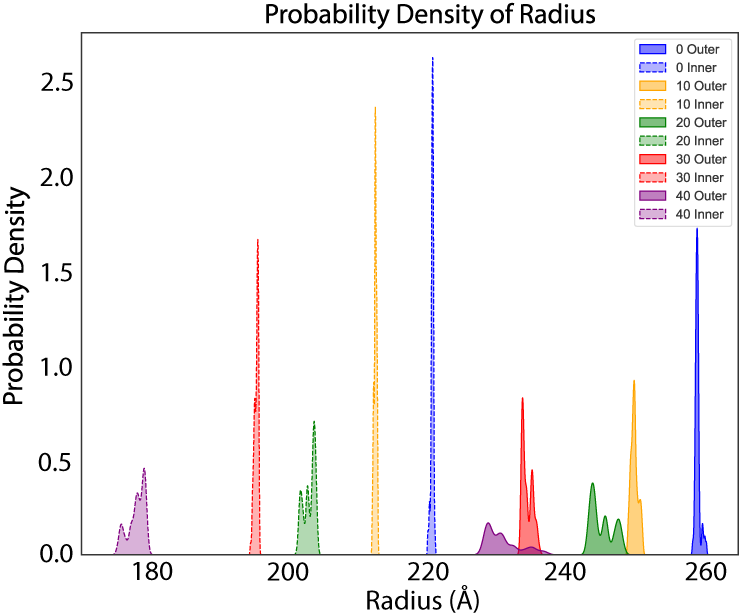
Cholesterol-dependent radius distributions of spherical lipid bilayers. Probability density distributions of the inner and outer radii for spherical DOPC/CHOL bilayers at different cholesterol concentrations. Increasing cholesterol fraction shifts both distributions toward smaller radii under fixed lipid counts, indicating composition-dependent vesicle relaxation. Broader distributions at higher cholesterol concentrations reflect increased variability associated with cholesterol-modulated lipid packing and membrane order within the CG framework.

In addition to shifts in the mean radius, higher cholesterol concentrations are associated with broader radius distributions. This increased dispersion reflects greater variability in vesicle relaxation across replicas and is attributed to the competing influences of cholesterol-induced ordering and curvature related packing stress in spherical geometries [43, 57]. Similar composition-dependent variability has been reported in previous simulations of cholesterol containing membranes.

Overall, the radius probability analysis indicates that cholesterol modulates vesicle geometry in a concentration dependent manner by altering lipid packing and membrane order. These trends should be interpreted as relative, model-dependent responses within a CG framework, rather than as quantitative predictions of experimentally measured vesicle sizes.

## 4. Conclusion

In this work, we used CG-MD simulations to investigate how cholesterol concentration and membrane curvature jointly influence the structural and dynamic behavior of DOPC-based lipid membranes. By systematically comparing planar bilayers and spherical vesicles across a controlled range of cholesterol contents, we isolated geometry-dependent trends that arise from cholesterol-mediated lipid packing and bilayer organization within a consistent MARTINI framework.

Our results reveal that cholesterol and curvature act as distinct and complementary control parameters governing membrane organization. Cholesterol primarily modulates intraleaflet structure, leading to increased membrane thickness, reduced solvent exposure, and enhanced lipid tail ordering across both geometries. In contrast, membrane curvature governs interleaflet coupling and structural heterogeneity, giving rise to increased interdigitation, elevated baseline ordering, and greater variability in membrane properties in vesicles relative to planar systems. These findings demonstrate that key structural descriptors derived from planar bilayers cannot be directly transferred to curved membranes without accounting for geometric constraints.

A key dynamic finding of this study is the non-monotonic dependence of apparent DOPC transbilayer exchange on cholesterol concentration in spherical bilayers, with a maximum observed near 20% cholesterol. This behavior does not indicate a universal or application-level optimal cholesterol fraction, nor does it represent a quantitative prediction of experimental flip-flop rates. Instead, it reflects a crossover regime specific to the simulated vesicle geometry and CG representation, in which curvature-induced leaflet stress and cholesterol driven ordering effects are most closely balanced.

Taken together, these results provide a coherent molecular level picture of how cholesterol and membrane curvature interact to shape lipid organization, ordering, and apparent transbilayer dynamics in simplified DOPC/cholesterol membranes. While the findings offer mechanistic insight within a CG framework, extension to experimentally relevant systems will require validation through atomistic simulations and experimental measurements, as well as exploration of additional lipid compositions, vesicle sizes, and force-field parameterizations.

## Author Contributions

E.K. conducted simulations, analyzed data, and wrote the manuscript. M.D.M. contributed to methodology. E.K. assisted with data analysis and molecular dynamics. M.M. designed the research, supervised the project, and edited the manuscript.

## Data Availability

All input files and analysis scripts necessary to reproduce the simulations and analyses reported in this study are publicly available at: https://github.com/bslgroup/liposome

## Acknowledgments

This research was supported by the NIH (R35GM147423), NSF (CHE 1945465), and the Arkansas Biosciences Institute. Computational resources were provided by the Texas Advanced Computing Center (TACC) at the University of Texas at Austin (Frontera) through LRAC allocation CHE21003. The work also used Stampede at TACC, Bridges-2 at the Pittsburgh Supercomputing Center, and Exapnse at San Diego Supercomputing Center through allocation MCB150129 from the Advanced Cyberinfrastructure Co-ordination Ecosystem: Services & Support (ACCESS) program. Additional computational support came from the Arkansas High-Performance Computing Center, funded by multiple NSF grants and the Arkansas Economic Development Commission.

