## Supporting Information for "Interplay Between Cholesterol Concentration and Membrane Curvature in Liposomes Revealed by Molecular Dynamics Simulations"

### 1 Cholesterol Structural Organization

Cholesterol (Fig. S1) was modeled using the MARTINI CG force field [1]. Its amphipathic architecture places the hydroxyl group near the headgroup region while the rigid sterol rings align with lipid acyl chains, allowing cholesterol to modulate packing, increase tail ordering, and enhance membrane mechanical stability [2, 3].

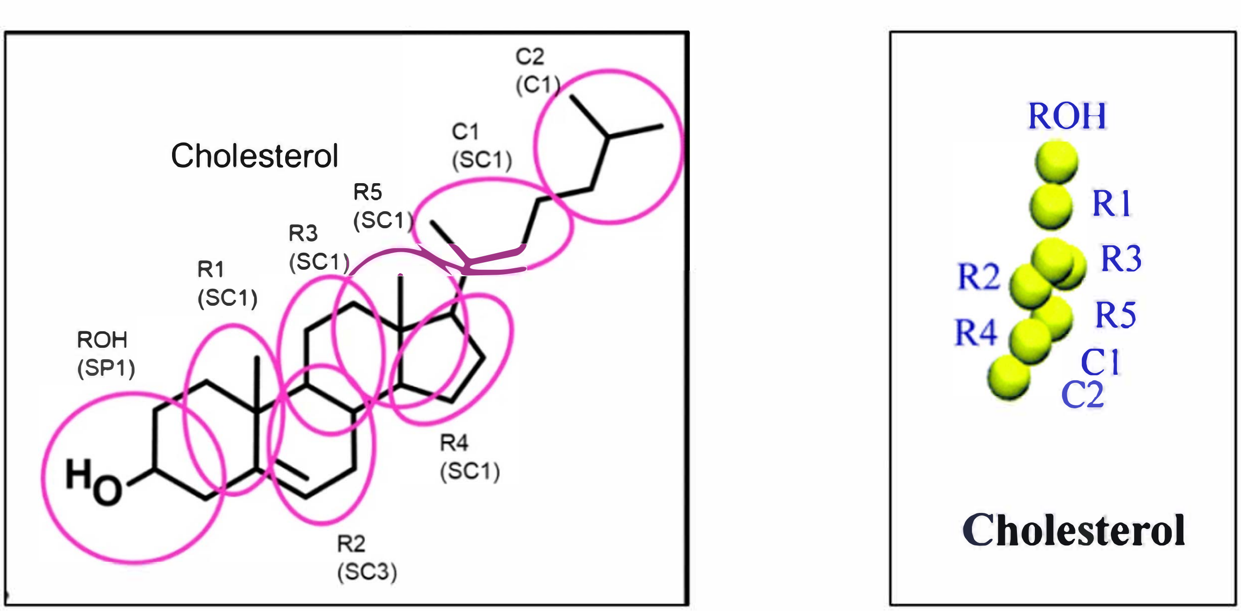


Figure S1: CG representation and structural orientation of cholesterol within the membrane.

#### 2 CG representation of DOPC

DOPC (Fig. S2) is a zwitterionic phospholipid that forms liquid-disordered, fluid bilayers at physiological temperature due to its unsaturated acyl chains [4, 5]. In the MARTINI representation, DOPC preserves key headgroup polarity and tail flexibility, enabling efficient sampling of sterol–lipid interactions and membrane structural responses [6].

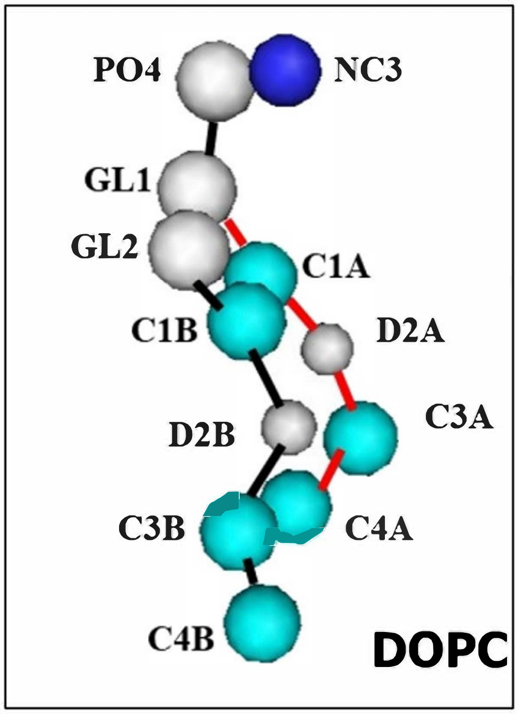


Figure S2: CG representation of DOPC used in the simulations.

#### 3 Planar vs Spherical Comparison at 20% Cholesterol

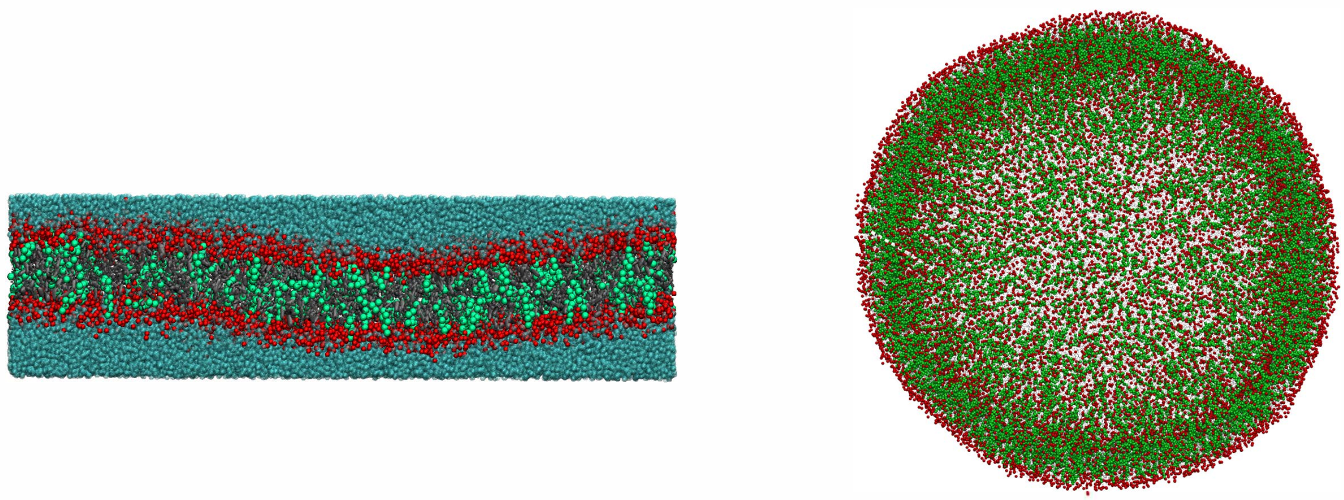

Snapshots of the Planar bilayer at 310 K with 20% cholesterol at the upper and lower leaflets.

Snapshots of the Spherical bilayer at 310 K with 20% cholesterol.

Figure S3: Side-by-side structural comparison of planar and spherical bilayers at 20% cholesterol concentration (310 K).

### 4 Structural Snapshots of Planar and Spherical Membranes

Representative configurations of the planar (Fig. S4) and spherical (Fig. S5) systems at 310 K are shown for the full cholesterol concentration series. These snapshots provide a qualitative visualization of membrane morphology and cholesterol distribution as a function of sterol content and geometry.

#### 4.1 Planar bilayers (0–40% cholesterol)

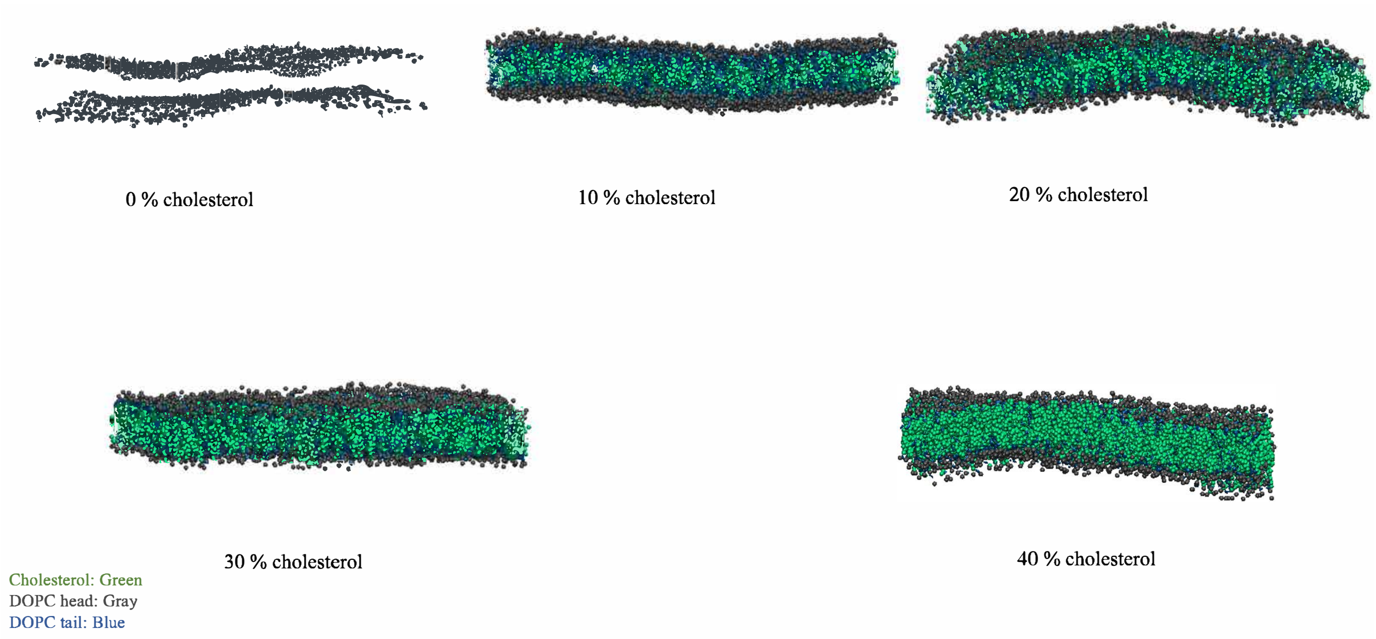


Figure S4: Representative snapshots of planar bilayers at 310 K for 0, 10, 20, 30, and 40% cholesterol in both leaflets.

#### 4.2 Spherical bilayers (0–40% cholesterol)

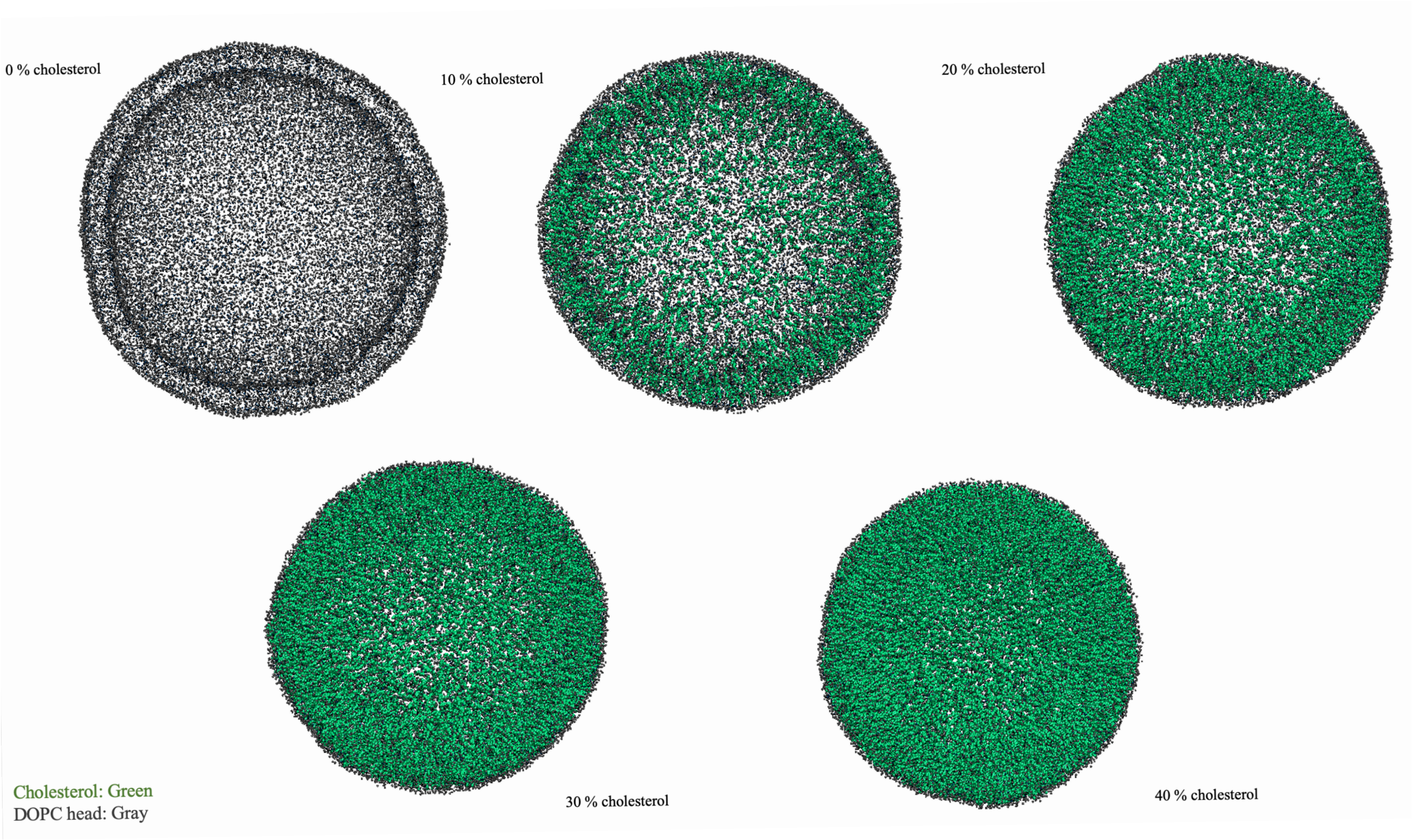


Figure S5: Representative snapshots of spherical bilayers at 310 K for 0, 10, 20, 30, and 40% cholesterol in both leaflets.

### 5 Segmental Order Parameter ($S_{CD}$)

Lipid tail orientational order was quantified using the segmental (deuterium) order parameter, defined as

$$S_{CD} = \frac{1}{2}\left(3\cos^2\theta - 1\right), \tag{1}$$

where $\theta$ is the angle between a lipid acyl chain bond vector and the membrane normal [7]. The order parameter provides a measure of the average orientational alignment of lipid tail segments relative to the bilayer normal. Higher $S_{CD}$ values indicate increased orientational ordering and reduced conformational flexibility of the acyl chains, corresponding to a more rigid and tightly packed membrane environment [8].

For each lipid, bond vectors along the acyl chains were constructed between consecutive coarse-grained tail beads. The angle $\theta$ was computed for each segment at every analyzed frame. $S_{CD}$ values were calculated separately for the first (sn-1) and second (sn-2) acyl chains and subsequently averaged over all lipids and over equilibrated production frames to obtain statistically converged profiles.

##### Definition of the Membrane Normal

For planar bilayers (Fig. S6), the membrane normal was defined along the global bilayer normal (z-axis) of the simulation box. Because planar systems maintain translational symmetry in the lateral ($x$–$y$) plane, this global normal provides a consistent reference direction for all lipids.

For spherical bilayers (Fig. S7), lipid orientations were evaluated relative to curvature-aware local membrane normals. Specifically, for each lipid, the local normal direction was defined as the radial vector connecting the vesicle center of mass to the corresponding lipid headgroup position. This curvature-dependent definition ensures that tail alignment is evaluated relative to the local membrane surface rather than a fixed Cartesian axis, thereby accounting for geometric curvature effects [9].

##### Cholesterol-Dependent Ordering

In planar bilayers (Fig. S6), $S_{CD}$ values increase systematically with cholesterol concentration for both sn-1 and sn-2 chains. This trend reflects cholesterol's well-established ordering and condensing effect in phosphatidylcholine membranes. Cholesterol inserts between phospholipid molecules, restricts acyl chain conformational freedom, and promotes trans conformations, leading to enhanced orientational alignment along the bilayer normal [2, 3].

Spherical bilayers (Fig. S7) exhibit the same qualitative cholesterol-dependent increase in $S_{CD}$. However, modest replicate-to-replicate variability is observed relative to planar systems. This increased dispersion arises from curvature-induced geometric heterogeneity: curved membranes impose spatially varying packing constraints across the vesicle surface, which can lead to local differences in acyl chain orientation and packing stress [10]. Importantly, the monotonic increase of $S_{CD}$ with cholesterol concentration is preserved across both geometries, indicating that cholesterol's ordering influence remains robust in curved membranes.

Overall, the $S_{CD}$ analysis demonstrates that cholesterol enhances lipid tail ordering in both planar and spherical bilayers within the MARTINI CG framework. Membrane curvature does not alter the direction of cholesterol-dependent ordering trends but modestly increases the variability of order parameter profiles due to geometric constraints. Because orientational relaxation is accelerated in CG models relative to atomistic simulations and experiments, the reported $S_{CD}$ values should be interpreted as relative structural descriptors rather than direct quantitative predictions of experimental deuterium order parameters.

#### 5.1 Planar bilayers: $S_{CD}$ profiles across cholesterol concentration

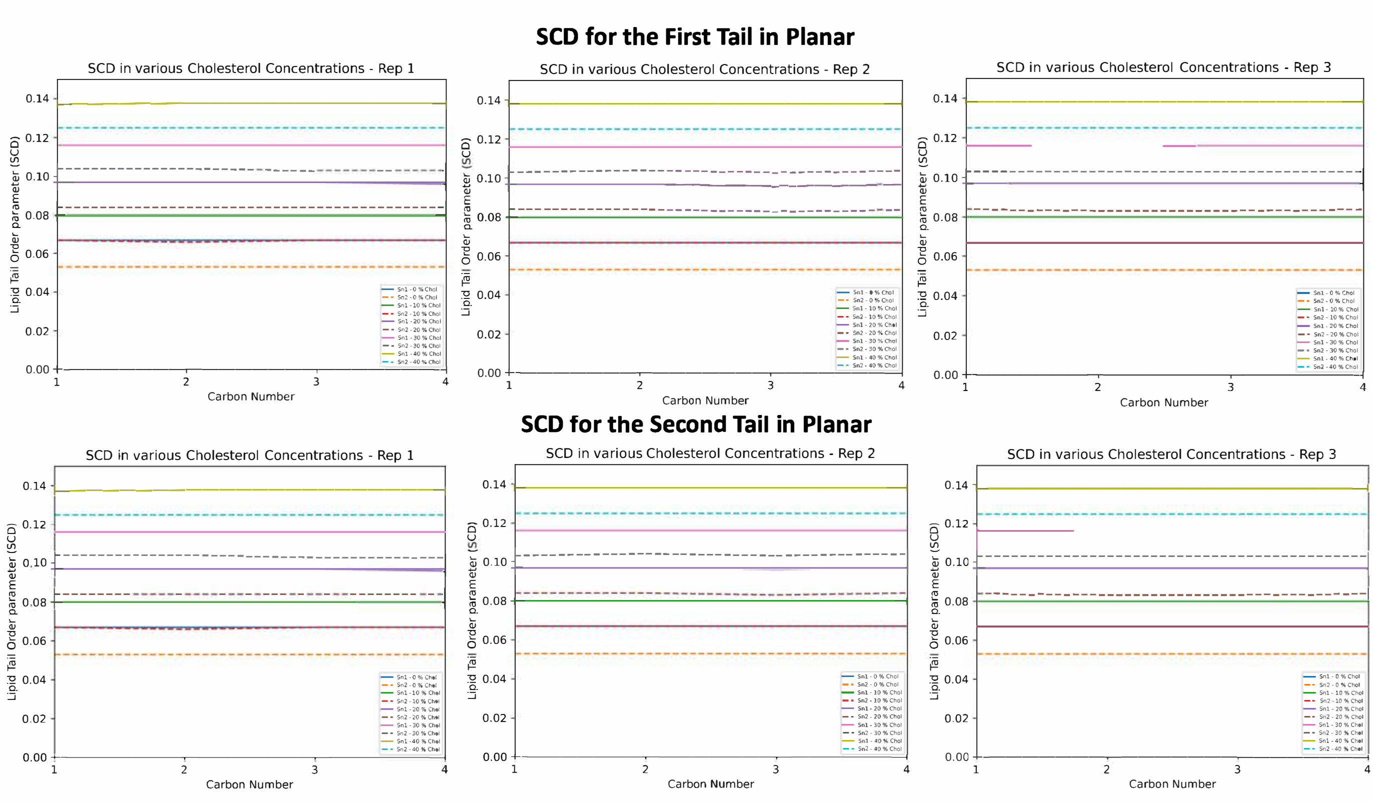


Figure S6: Segmental order parameter profiles for planar bilayers across cholesterol concentrations (three independent replicates).

#### 5.2 Spherical bilayers: $S_{CD}$ profiles across cholesterol concentration

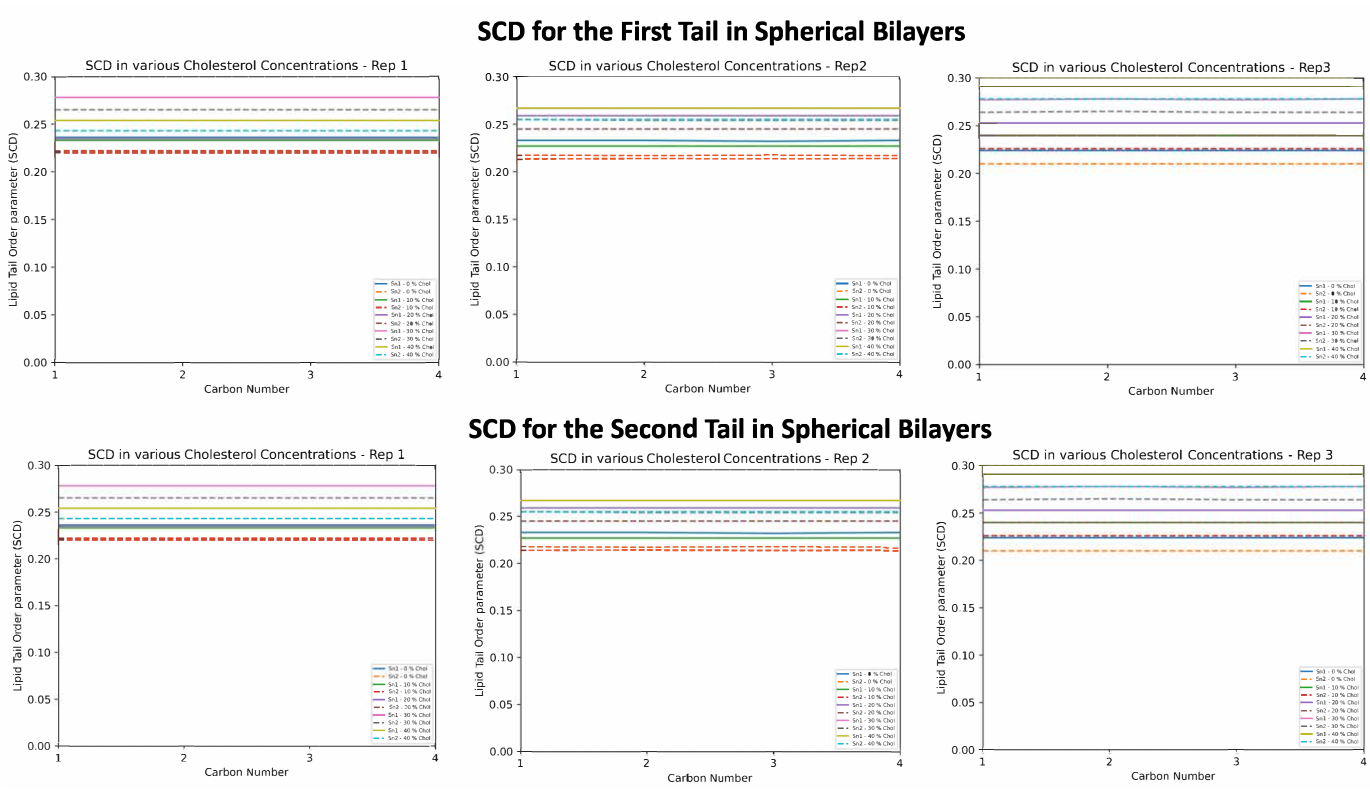


Figure S7: Segmental order parameter profiles for spherical bilayers across cholesterol concentrations (three independent replicates).

### 6 Supplementary Analysis of Area per Lipid

The average area per lipid (APL) was calculated to quantify cholesterol-dependent changes in lateral molecular packing as a function of membrane geometry. APL provides a direct structural descriptor of lipid condensation and packing efficiency and has been widely used in both experimental and computational studies of membrane organization [11, 12].

For planar bilayers, APL was computed using two-dimensional Voronoi tessellation applied to the projected $x$–$y$ coordinates of lipid headgroup beads, following established methodologies for flat membranes [11]. In this approach, the membrane plane is partitioned into polygonal regions, each associated with a single lipid molecule. The area of each Voronoi cell represents the local lateral packing area of that lipid. APL values were obtained by averaging over all lipids and equilibrated trajectory frames, yielding a time-averaged measure of molecular packing.

For spherical bilayers, direct planar tessellation is not applicable due to membrane curvature. To account for geometric effects, lipid headgroup positions were mapped onto the vesicle surface, and an effective surface area per lipid was estimated by dividing the total vesicle surface area by the number of lipids in each leaflet [9]. This procedure yields a global, curvature-aware APL metric that enables consistent comparison between planar and spherical systems without imposing planar tiling assumptions. Accordingly, the reported APL values represent global averages rather than spatially resolved local packing variations.

Using this definition, both planar and spherical bilayers exhibit a monotonic decrease in APL for DOPC and cholesterol molecules with increasing cholesterol concentration (Fig. S8). In planar bilayers, the average area per DOPC decreases systematically as cholesterol fraction increases, consistent with cholesterol's well-established condensing effect in phosphatidylcholine membranes [2, 3]. Sterol–lipid interactions promote tighter lateral packing, restrict acyl chain conformational freedom, and reduce the effective molecular area of phospholipids. The average area per cholesterol molecule in planar bilayers also decreases gradually with cholesterol fraction, reflecting increasingly efficient sterol packing and enhanced sterol–sterol contacts at higher concentrations [13].

Spherical bilayers display the same qualitative cholesterol-dependent trends observed in planar systems. Both DOPC and cholesterol APL values decrease with increasing sterol content, demonstrating that cholesterol retains its condensing influence in curved geometries. However, spherical systems exhibit modestly larger replicate-to-replicate variability compared to planar membranes. This increased dispersion arises from curvature-induced packing heterogeneity: curved membranes impose spatially varying geometric constraints on lipid organization, leading to continuous differences in local molecular environments across the vesicle surface [9]. Importantly, this variability reflects geometric heterogeneity rather than the formation of discrete packing domains.

Overall, cholesterol induces a robust and monotonic reduction in average molecular area for both DOPC and cholesterol across planar and spherical bilayers. Membrane curvature does not alter the direction of cholesterol-dependent packing trends but increases the variability of area estimates due to geometric constraints. Within the MARTINI CG framework, these results demonstrate that cholesterol's condensing effect on lipid packing is preserved across membrane geometries, while curvature primarily modulates dispersion rather than the qualitative behavior of lipid area measurements.

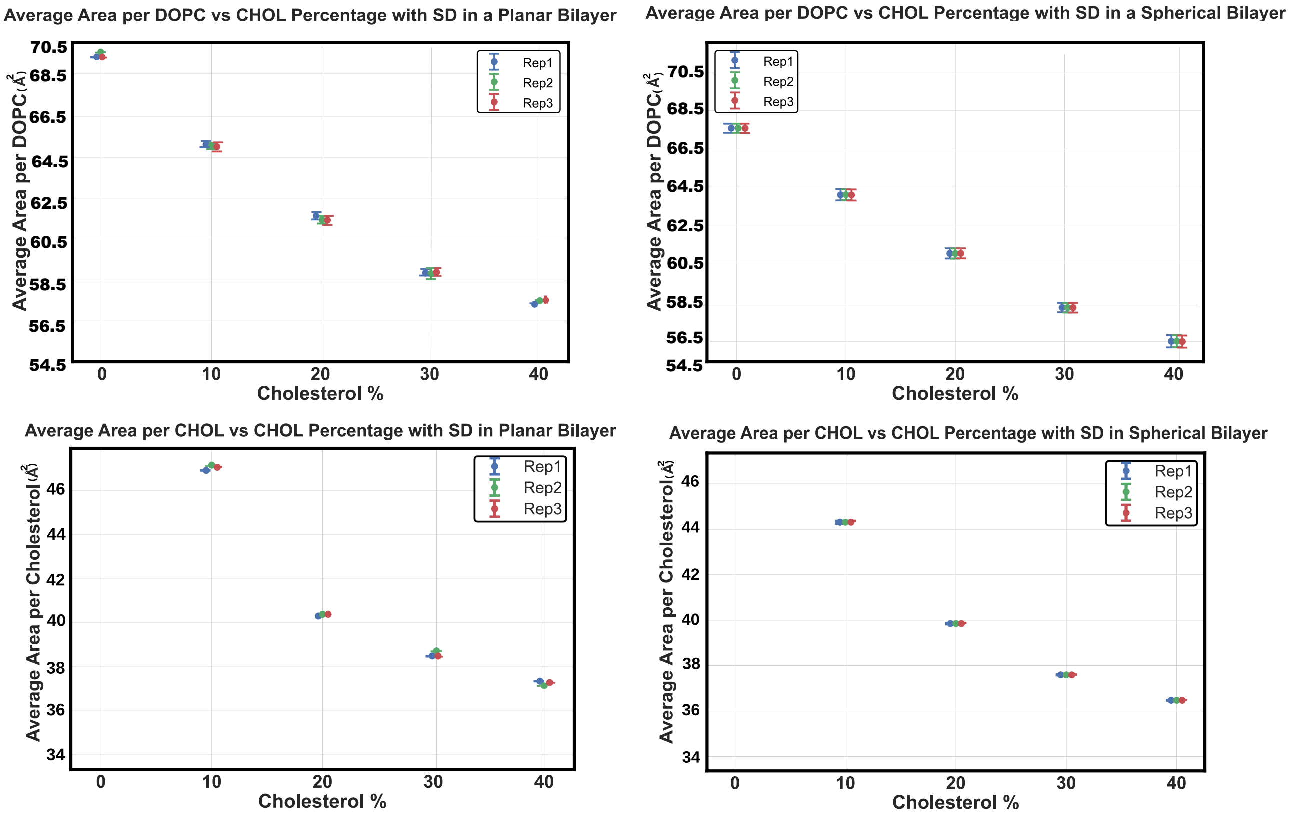


Figure S8: Cholesterol-dependent variation in average area per lipid for planar and spherical membranes. Both DOPC and cholesterol exhibit a monotonic decrease in APL with increasing sterol concentration. Spherical bilayers show modestly larger variability due to curvature-induced packing heterogeneity.

### 7 Curvature-Induced Leaflet Asymmetry

To complement the transbilayer exchange analysis described in the main text, lipid distributions between opposing membrane leaflets were quantified to assess curvature-associated redistribution as a function of cholesterol concentration and membrane geometry. Lipids were assigned to leaflets on a frame-by-frame basis using the same geometric criteria employed for flip-flop analysis. Specifically, lipid headgroup bead positions were referenced to the instantaneous membrane midplane [14].

For planar bilayers, the membrane midplane was defined relative to the global bilayer normal (z-axis), and lipids were classified as belonging to the upper or lower leaflet according to their position relative to this plane. For spherical bilayers, the midplane was defined with respect to the vesicle center of mass, and lipids were classified as inner or outer leaflet components based on their radial distance from the vesicle center.

For each trajectory, the number of DOPC and cholesterol molecules in each leaflet was computed at every analyzed frame and averaged over the equilibrated production segment of the simulation. This procedure yields a time-averaged compositional measure of lipid distribution between leaflets within the coarse-grained framework.

In planar bilayers (Fig. S9), leaflet populations remain symmetric across all cholesterol concentrations. The close overlap between DOPC and cholesterol fractions in the upper and lower leaflets confirms that the simulation setup and leaflet assignment protocol do not introduce artificial compositional bias in flat membranes. This symmetry therefore serves as an internal validation of the analysis methodology.

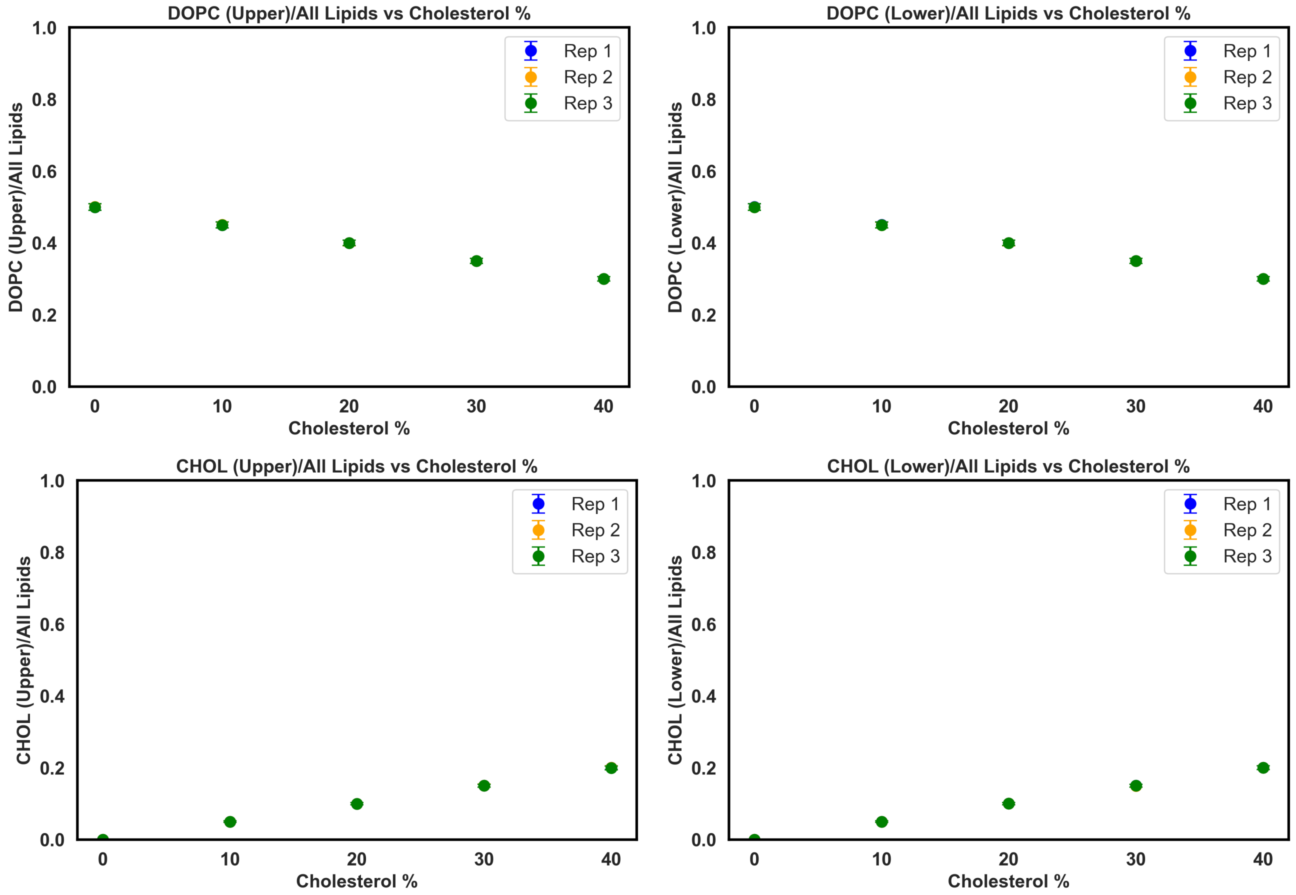


Figure S9: Leaflet lipid distribution in planar bilayers. Fractions of DOPC and cholesterol in upper and lower leaflets as a function of cholesterol concentration. The near-identical leaflet populations confirm the absence of curvature-induced redistribution in flat membranes.

In spherical bilayers (Fig. S10), modest differences between inner and outer leaflet populations are observed. Unlike planar systems, curved membranes inherently impose geometric asymmetry: the outer leaflet possesses a larger surface area and experiences lateral expansion, whereas the inner leaflet is subjected to geometric compression. These curvature-induced differences in available surface area and packing constraints can influence lipid accommodation during equilibration. The observed leaflet-dependent variations are therefore interpreted as curvature-associated redistribution arising from differential packing constraints rather than persistent thermodynamic compositional asymmetry [9, 15].

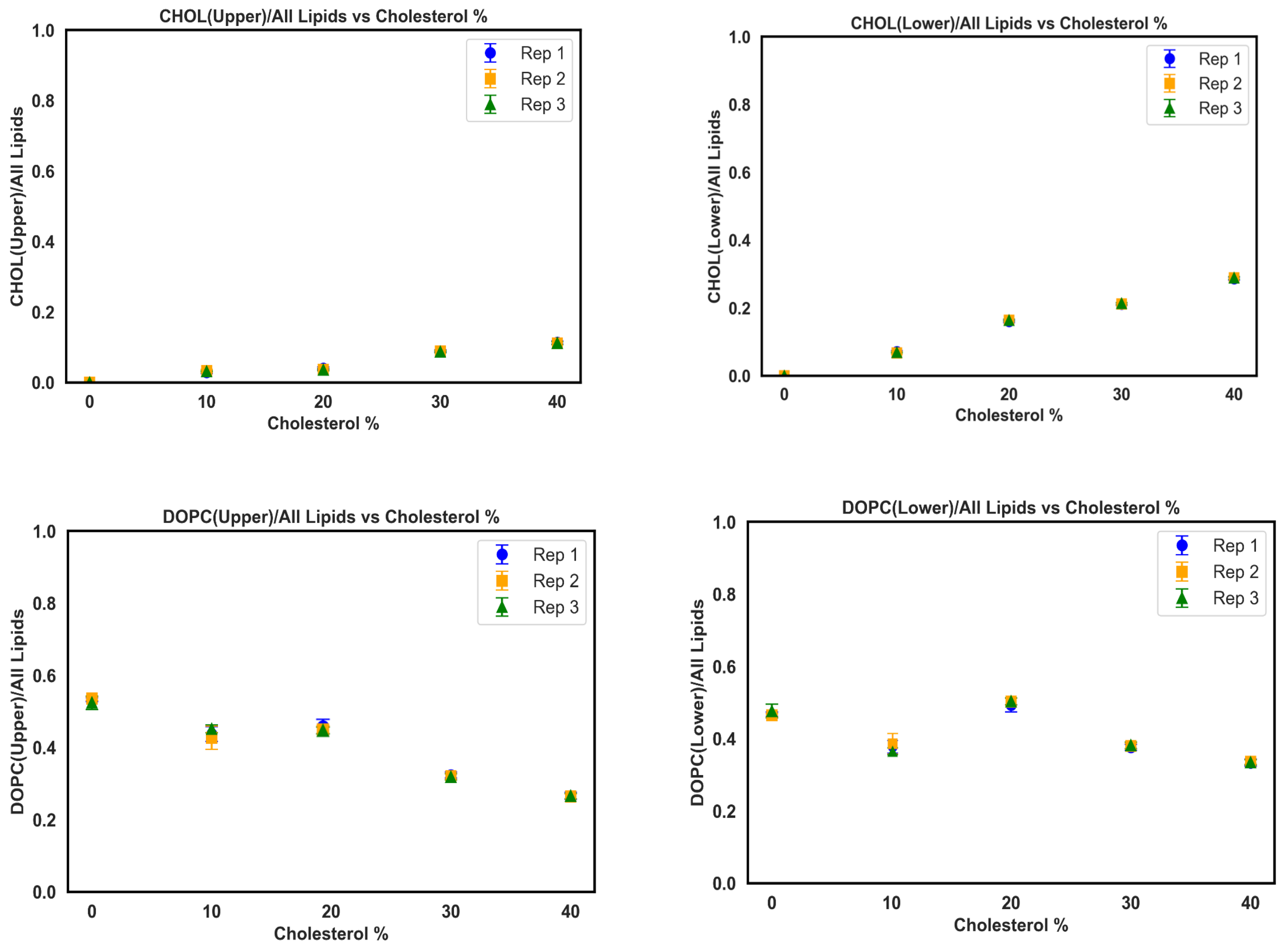


Figure S10: Leaflet lipid distribution in spherical bilayers. Inner and outer leaflet lipid fractions as a function of cholesterol concentration. Modest leaflet-dependent differences reflect curvature-induced packing asymmetry between compressed inner and expanded outer leaflets.

Because lipid diffusion and relaxation processes are accelerated in the MARTINI CG representation relative to atomistic simulations and experimental timescales [6, 16], the reported leaflet population differences should be interpreted as qualitative, model-dependent descriptors of curvature-driven lipid redistribution rather than quantitative predictors of equilibrium leaflet compositions in biological membranes.
